# Mice Lacking FXR Are Susceptible to Liver Ischemia-Reperfusion Injury

**DOI:** 10.1101/739839

**Authors:** Yuxin Li, Rui Xu, Prahlad K. Rao, Charles K Gomes, E. Richard Moran, Michelle Puchowicz, Eugene B. Chang, Deng Ping Yin, Joseph F. Pierre

## Abstract

Activation of bile acid (BA) receptor, farnesoid X receptor (FXR) has been shown to inhibit inflammatory responses and improve tissue ischemia-reperfusion injury (IRI). This study investigated the effect of FXR deficiency on liver IRI, using a liver warm IRI mouse model. We demonstrate that liver IRI resulted in decreased FXR expression in the liver of WT mice. FXR^-/-^mice displayed greater liver damage and inflammatory responses than WT mice, characterized by significant increases in liver weight, serum AST and ALT, hepatocyte apoptosis and liver inflammatory cytokines. Liver IRI increased expression of X box binding protein 1 (XBP1) and FGF21 in WT liver, but not in FXR^-/-^ liver, which conversely increased CHOP expression, suggesting a loss of ER stress protection in the absence of FXR. FXR deficiency increased circulating total BAs and altered BA composition with reduced TUDCA and hepatic BA synthesis markers. FXR deficiency also reshaped gut microbiota composition with increased Bacteroidetes and Proteobacteria and decreased Firmicutes. Curiously, Bacteroidetes were positively and Firmicutes were negatively correlated with serum ALT levels. Administration of FXR agonist CDCA inhibited NF-*κ*B activity and TNFα expression *in vitro* and improved liver IRI *in vivo*. Our findings demonstrate that FXR signaling plays an important role in the modulation of liver IRI.

## Introduction

Recent improvements in surgical techniques, liver preservation and immunosuppression continue to improve liver operations and increase survival of liver grafts in transplantation (1, 2). However, liver ischemia and reperfusion injury (IRI) remains an important problem in the clinical scenario of liver surgery (3). In major liver resection, a primary side effect is liver warm IRI, i.e. ischemia of remnant liver following temporary vascular occlusion, and additional reperfusion injury added to the damage sustained during ischemia (4). In liver transplantation, liver warm and cold IRI may occur *in situ* during recipient surgery or donor liver harvest, which may cause cellular injury, organ dysfunction, or even complete graft failure (5).

Tissue ischemia and reperfusion initiates a defensive process known as the unfolded protein response (UPR) for adaptation and safeguard of cell survival. Activation of the inositol-requiring enzyme (IRE)/X-box binding protein 1 (XBP1) pathway in response to the endoplasmic reticulum (ER) stress protects hepatocytes from apoptosis. However, sustained activation of ER triggers proapoptotic signals via C/EBP homologous protein (CHOP), which is responsible for cellular dysfunction (6, 7). Evidence for the role of BAs and their receptors in the regulation of inflammation and tissue IRI already exists in experimental and animal settings (8, 9). Activation of the BA membrane receptor, G protein-coupled BA receptor 1 (TGR5), by TGR5 agonists inhibits inflammatory responses and attenuates liver IRI by suppressing the Toll like receptor 4 (TLR4)-NF-*κ*B mediated pathway (10). Recent experimental data suggest that activation of BA nuclear receptor, farnesoid X receptor (FXR), stimulates the IRE/XBP1 pathway and may be protective during liver injury (11). Similarly, administration of the FXR-agonist obeticholic acid (OCA) improves survival in a rodent model of intestinal IRI, through gut barrier preservation and mitigated inflammation (12). However, in the setting of liver IRI, the role and fundamental mechanism of FXR signaling in the modulation of ER stress and inflammatory responses remain unexplored.

There is growing evidence of bidirectional interactions between BAs and the gut microbiota, i.e. BAs in the distal intestine influence the composition of the gut microbiota that in turn modify primary BAs to generate secondary BAs (13). FXR deficiency significantly reduces the abundance of Firmicutes and increased Bacteroidetes (14). Dysbiosis directly impairs intestinal barrier function and increases gut permeability. Bacteria and their components, such as lipopolysaccharide (LPS), pass through the intestinal barrier to the liver, causing inflammatory responses. Modulation of the gut microbiota may inhibit inflammatory responses and represent a novel therapeutic approach for the prevention of tissue IRI (15).

In this study, we examined FXR deficiency in the aggravation of liver IRI using a mouse model of liver warm IRI. We hypothesized that a lack of FXR exaggerates hepatocyte ER stress and apoptosis during liver IRI. FXR deficiency may alter BA and gut microbiota homeostasis, which may also contribute to the pathogenesis of liver IRI. Our results demonstrate that mice lacking FXR are more susceptible to liver IRI than wild type (WT) mice.

## Materials and Methods

### Mice and surgical procedures

FXR knockout (FXR^-/-^, B6.129X1-*Nr1h4tm1Gonz*/J) mice (8-weeks of age) were purchased from Jackson Laboratory. Male homozygous FXR^-/-^and FXR wild type (WT) littermate mice (5 mice per group) were used. All mice were housed in a specific pathogen-free, temperature-controlled and 12-hour light-dark cycle environment in the Animal Resources Center (ARC) at the University of Chicago.

The liver warm IRI model was conducted under general anesthesia with isoflurane and equal O2 through a cone placed around the mouse’s nose. A small clip (Roboz, RS-5424) was used to interrupt arterial/portal venous blood supply to the median and left lateral lobes, in which 70% of liver blood flow was blocked (16). To test whether a lack of FXR exaggerates liver damage, after 90 minutes of ischemia, the clip was removed followed by 6 hours of reperfusion in FXR^-/-^mice compared to WT mice. To test whether FXR activation improved liver IRI, FXR agonist chenodeoxycholic acid (CDCA) was administered to WT mice at a dose of 150 mg/kg (dissolved in 100μl 1% carboxymethylcellulose or CMC) by oral gavage daily for a week prior to IRI challenge. The same volume of CMC (Vehicle) was used in control mice. The animal’s vital signs were monitored throughout surgery, including respiratory rate, response to pedal or palpebral stimulus and assessment of spontaneous movements. All animals received humane care, according to the criteria outlined in the “Guide for the Care and Use of Laboratory Animals” prepared by the National Academy of Sciences and published by the NIH. All animal protocols were approved by the Institutional Animal Care and Use Committee (IACUC) at the University of Chicago.

### Serum alanine aminotransferase (ALT) and aspartate aminotransferase (AST) analysis

Blood samples were collected at 6 hours (WT vs. FXR^-/-^) or 24 hours (WT with and without FXR agonist CDCA) after ischemia and were centrifuged to obtain serum. ALT and AST were measured to assess the extent of hepatocyte damage using an automated chemical analyzer (Olympus Automated Chemistry Analyzer AU5400, Tokyo, Japan).

### Bile acid analysis

Total BAs and BA composition were analyzed as described in our previous reports (17, 18). Briefly, stock solutions of individual BAs and NDCA were prepared in methanol at a concentration of 5μg/mL. Calibration standards were prepared by adding individual bile acids at a concentration range of 12ng/mL to 1.5ug/mL to charcoal stripped human serum. 1.6μL of NDCA was added to 40μL of standards and samples. Deproteinization was carried out by adding 15X ice-cold methanol to 40μL of standards and samples. The supernatant was transferred to a new tube, evaporated under vacuum and dissolved in 100μL of 50% methanol. The tubes were centrifuged at 11,000xg for 1 min before transfer in to specific vials for injection in to an LC-MS/MS system. Data were acquired on AB Sciex triple quadrupole mass spectrometer in negative ion mode coupled to Shimadzu Nexera XR HPLC system. Chromatographic separation was carried out on Thermo Scientific Accucore XLC8 column (4μm, 100 x 3mm I.D.). Quantitation of bile acids was carried out on MultiQuant software v3.0.2 (AB Sciex). BAs, including CA, CDCA, DCA, LCA, TCA, TDCA, TCDCA, UDCA and TUDCA, were purchased from Sigma-Aldrich. bMCA, TbMCA and internal standard 23-nordeoxycholic acid (NDCA) was purchased from Toronto Research Chemicals (Toronto, Ontario, Canada).

### Cecal content microbiota analysis

Gut microbiota in the cecal content was analyzed as described in our previous report (17, 18). Briefly, primers specific for 16S rRNA V4-V5 region (Forward: 515F: 5’-GTGYCAGCMGCCGCGGTAA −3’ and Reverse: 806R: 5’-GGACTACHVGGGTWTCTAAT-3’) that contained Illumina 3’ adapter sequences, as well as a 12-bp barcode were used. Sequences were generated by an Illumina MiSeq DNA platform at Argonne National Laboratory and analyzed by the program Quantitative Insights Into Microbial Ecology (QIIME) (19). Operational Taxonomic Units (OTUs) were picked at 97% sequence identity using open reference OTU picking against the Greengenes database (20). OTUs generated in QIIME were then analyzed using linear discriminant analysis (LDA) effect size (LEfSe) where non-parametric factorial Kruskal-Wallis sum-rank testing (*p* < 0.05) identified significantly abundant taxa followed by unpaired Wilcoxon rank-sum test to determine LDA scores > 2 (21).

### Real-time RT-PCR for quantification of mRNA expression

Liver tissue was collected at the time of animal euthanization and immediately placed in Trizol reagent (Ambion, Austin, TX) (22, 23). Total RNA was reverse-transcribed to complementary DNA (cDNA) using the Transcriptor First Strand cDNA Synthesis Kit (Roche, Indianapolis, IN). RT-PCR amplification was consisted of an initial denaturation step (95°C for 10 min), 45 cycles of denaturation (95°C for 10s), annealing (55°C for 20s) and extension (60°C for 30s), followed by a final incubation at 55°C for 30s and cooling at 40°C for 30s. All measurements were normalized by the expression of GAPDH gene, considered as a stable housekeeping gene. Gene expression was determined using the delta-delta Ct method: 2-ΔΔCT (ΔΔCT = [Ct(target gene) – Ct(GAPDH)] tested - [Ct(target gene) -Ct(GAPDH)]control) and displayed as relative mRNA levels.

### Cell culture and tissue luciferase assay

To test whether the FXR agonist CDCA inhibited NF-*κ*B activity and TNFα expression in macrophages, bone marrow-derived macrophages (BMDMs) transfected with a luciferase gene under the control of the NF-*κ*B promoter (kindly provided by Dr. Wei Han, Vanderbilt University) (24) were incubated with lipopolysaccharide (LPS, 0.025μg/ml, Sigma-Aldrich, St. Louise, MO). BMDMs were maintained in DMEM (Invitrogen, Carlsbad, CA) supplemented with 10% FCS at 37°C in a 5% CO_2_ incubator for 4 hours. Luciferase activity in BMDMs was determined by the tissue luciferase assay using the Bradford Luciferase Reporter Assay kit (Promega, Madison, WI, USA) following the manufacturer’s instructions as described in our previous report (25). TNFα mRNA was analyzed by qRT-PCR as described above.

### Histological examination

Liver tissue was collected 6 hours following ischemia and subjected to H&E staining. Hepatocyte apoptosis was analyzed by TUNEL assay. TUNEL labeling was performed using a TUNEL kit (Abcam, Cambridge, UK) according to the manufacturer’s instructions. Quantitation of apoptotic cells was accomplished by calculating of the labeling index, which was defined as the ratio between the number of labeled cells and the total cells counted in triplicate field by two blinded investigators.

### Statistical analysis

Data from the current studies were analyzed by ANOVA tests (StatView 4.5, Abacus Concepts, Berkeley, CA) with Tukey post-hoc multiple comparisons, or by 2-tailed Student’s *t* test when appropriate, where *p*-value < 0.05 was considered significant. The results are presented as mean ± SEM.

## Results

### Mice lacking FXR were more susceptible to liver IRI

A lack of FXR increased TGR5 expression in naïve FXR^-/-^mice, indicating a compensation of TGR5 for the deficiency of FXR. Liver IRI reduced hepatic expression of FXR in WT mice, and conversely, the same procedure increased expression of TGR5 in WT and FXR^-/-^mice (*Fig. 1, A and B*). FXR^-/-^mice also revealed an increase in liver weight (liver/body weight) in untreated FXR^-/-^(FXR^-/-^Control) mice, compared with untreated WT (WT Control) mice. Liver IRI further increased liver weight (Fig. 1*C*) and aggravated liver injury in the deficiency of FXR evidenced by increased circulating AST (Fig. 1*D*) and ALT (Fig. 1*E*) levels in FXR^-/-^IRI mice, compared with WT mice. The gross examination suggested evidence of liver steatosis and cholestasis in FXR^-/-^controls and FXR^-/-^IRI mice, which was confirmed histologically (H&E) (Fig. 1*F*), and suggested possible steatosis, which was previously identified as a susceptibility factor under IRI (26).

**Fig. 1.**
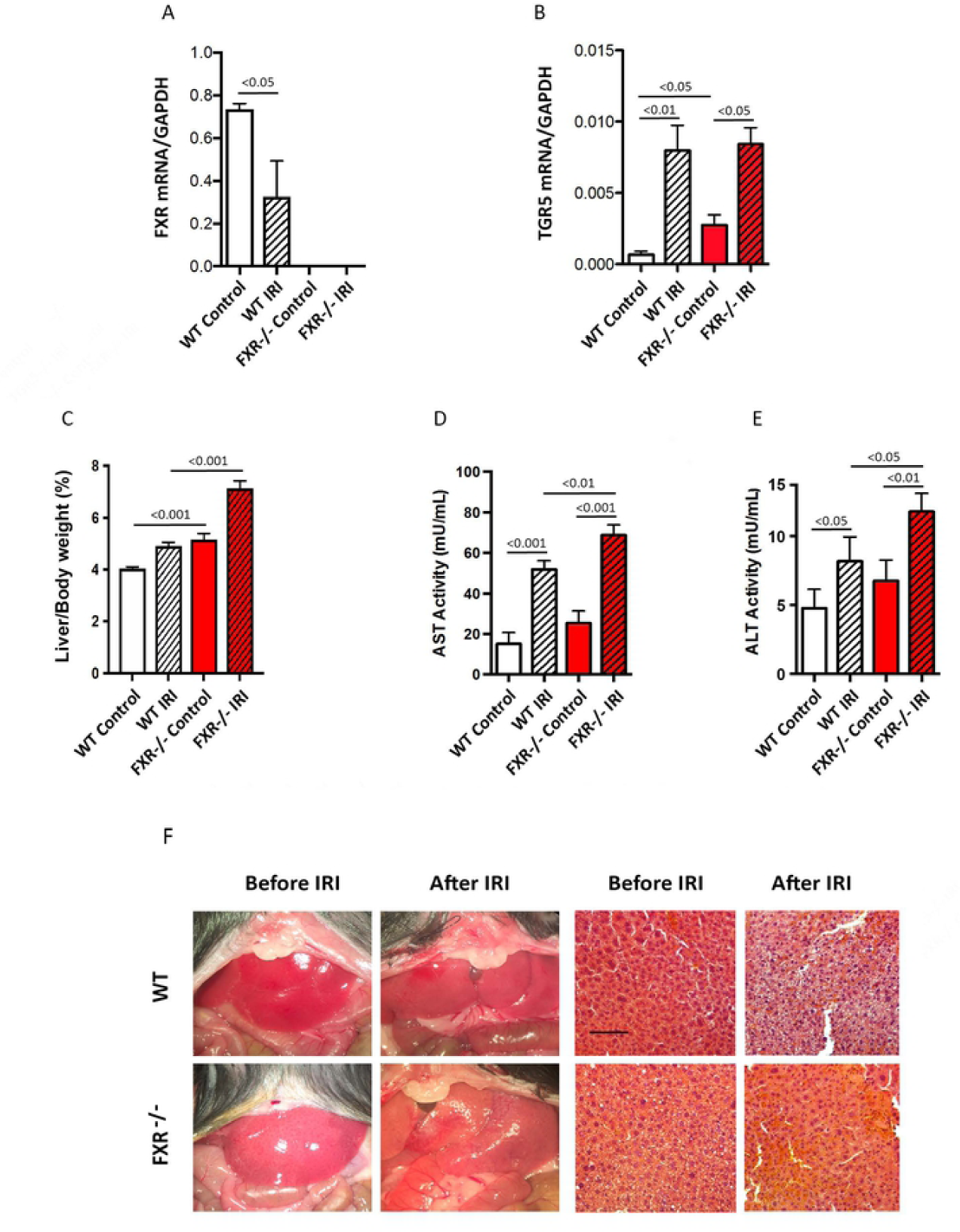
The lack of FXR was more susceptible to liver IRI (n = 5 per group). (A) Liver IRI decreased FXR expression in WT mice. WT IRI vs. WT Control, *p* < 0.05. (B) Liver IRI increased TGR5 expression in WT and FXR^-/-^mice. FXR^-/-^IRI vs. FXR^-/-^Control, *p* < 0.05; WT IRI vs. WT Control, *p* < 0.01, and FXR^-/-^Control vs. WT Control, *p* < 0.05. (C) Liver IRI increased liver weight in FXR^-/-^mice. FXR^-/-^Control vs. WT Control *p* < 0.01; FXR^-/-^IRI vs. WT IRI, *p* < 0.001. (D) The lack of FXR resulted in increase of serum AST. WT IRI vs. WT Control, *p* < 0.001; FXR^-/-^ IRI vs. FXR^-/-^Control, *p* < 0.001, and FXR^-/-^IRI vs. WT IRI, *p* < 0.01. (E) The lack of FXR resulted in increase of serum ALT. WT IRI vs. WT Control, *p* < 0.05; FXR^-/-^IRI vs. FXR^-/-^Control, *p* < 0.01, and FXR^-/-^IRI vs. WT IRI, *p* < 0.05. (F) The lack of FXR aggravated liver damage. Images for left panels are WT and FXR^-/-^livers, and right two panels showed histological changes (H&E staining) before and after IRI.

A lack of FXR resulted in increased TNFα, IL-1β and IL-6 transcript in FXR^-/-^control mice (FXR^-/-^Control vs. WT Control, *p* < 0.05, Fig. 2, *A to C*). As expected, liver IRI also increased TNFα, IL-1β and IL-6 transcripts in both WT and FXR^-/-^mice (*p* < 0.01), and TNFα was significantly increased in FXR^-/-^IRI mice, compared to WT IRI mice (Fig. 2*A*). Superoxide dismutase 2 (SOD2) binds to the superoxide to improve tissue injury. Reduced SOD2 expression was observed in WT mice following IRI, but more dramatically in FXR^-/-^mice (*p* = 0.05, Fig. 2*D*).

**Fig. 2.**
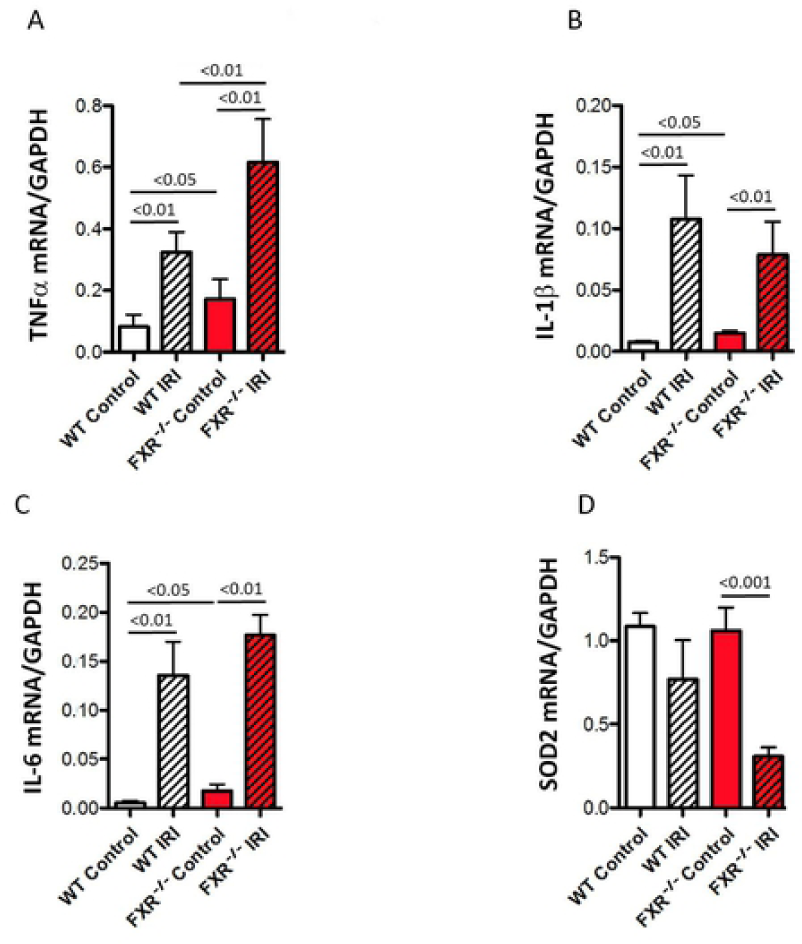
The lack of FXR increased expression of inflammatory cytokines (n = 5 per group). (A) Liver IRI and lack of FXR increased TNFα mRNA. WT Control vs. WT IRI, *p* < 0.01; WT Control vs. FXR^-/-^Control, *p* < 0.05; FXR^-/-^IRI vs. FXR^-/-^Control, *p* < 0.01 and FXR^-/-^IRI vs. WT IRI, *p* < 0.01. (B) Liver IRI and lack of FXR increased IL-1β expression. WT Control vs. FXR^-/-^Control, *p* < 0.05; WT Control vs. WT IRI, *p* < 0.01, and FXR^-/-^Control vs. FXR^-/-^IRI, *p* < 0.01. (C) Liver IRI and lack of FXR increased IL-6 expression. WT Control vs. WT IRI, *p* < 0.01; WT Control vs. FXR^-/-^Control, *p* < 0.05, and FXR^-/-^Control vs. FXR^-/-^IRI, *p* < 0.01. (D) The lack of FXR decreased SOD expression. FXR^-/-^IRI vs. FXR^-/-^Control, *p* < 0.01.

### FXR deficiency triggered hepatocyte apoptosis via activation of ER stress

ER stress activates a number of proteins that straddle ER membranes. Activated IRE1α functions as an endoribonuclease splicing a 26 base pair intron from XBP1 mRNA. Spliced XBP1 (sXBP1) mRNA is translated into a stable and active UPR transcription factor. Therefore, measuring XBP-1 splicing represents a reliable indirect method of determining IRE1α activation (27). Liver IRI promoted expression of total XBP1 (TXBP1, Fig. 3*A*), sXBP1 (Fig. 3*B*) and unconventional splicing XBP1 (usXBP1, Fig. 3*C*) in WT mice. However, mice deficient in FXR showed no change in these genes (Fig. 3, *A to C*). Liver IRI decreased expression of ATF4 and GFP94 in the absence of FXR (WT IRI vs. FXR^-/-^IR, *p* < 0.05, *Fig. 3, D and E*). Furthermore, FXR deficiency resulted in decreased ERAD-enhancing α-mannosidase-like protein (EDEM, Fig. 3*F*), ultimately resulting in hepatocyte susceptibility to apoptosis. Consistently, we observed increased CHOP with FXR deficiency (Fig. 3*G*). TUNEL tests showed increased apoptotic cells in untreated FXR^-/-^control mice, but without statistical difference. Liver IRI increased apoptotic cells in all groups; however, apoptotic cells were significantly increased in FXR^-/-^mice compared with WT IRI mice (Fig. 3*H*).

**Fig. 3.**
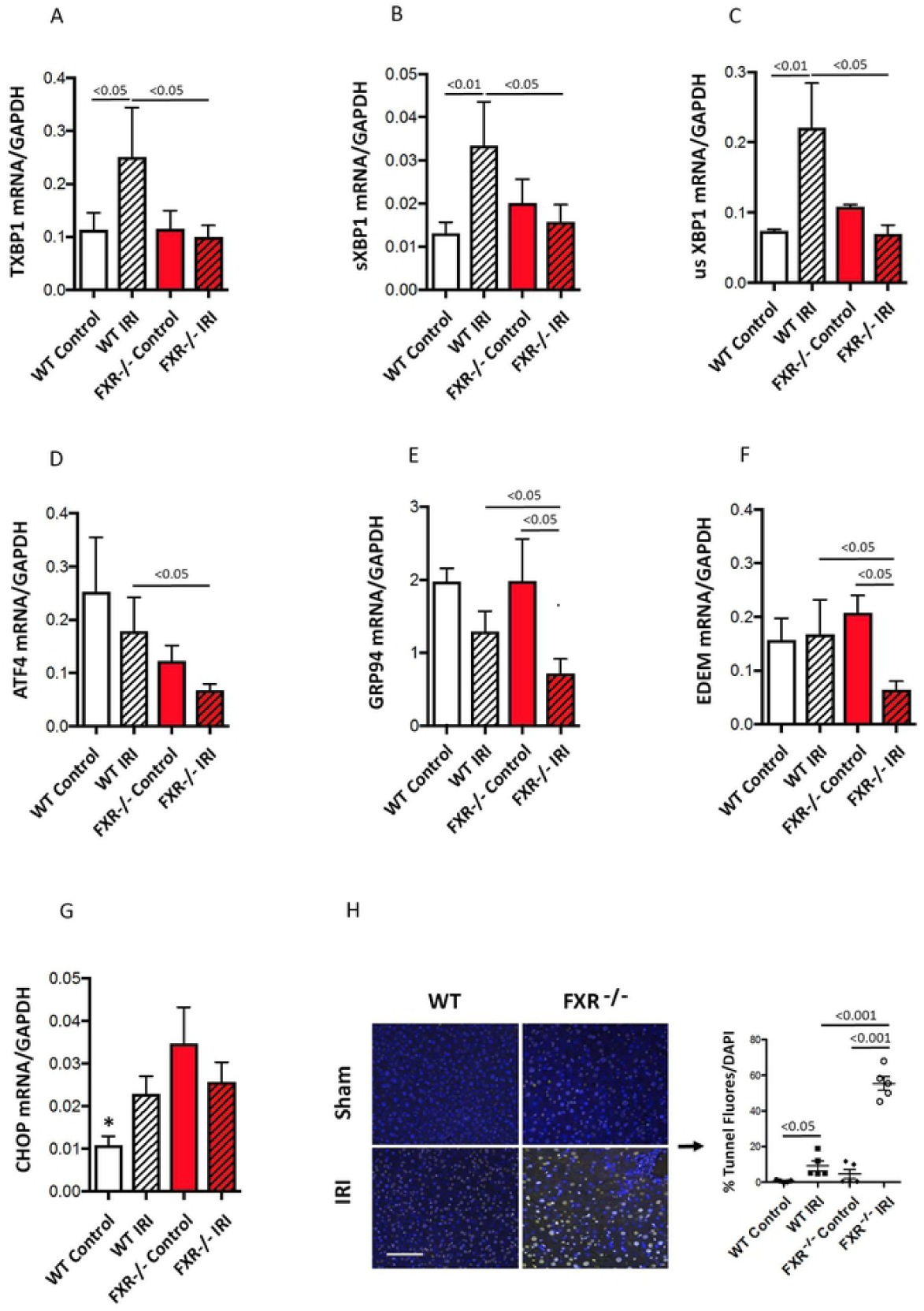
The lack of FXR altered gene expression of ER stress markers (n = 5 per group). (A) FXR deficiency failed to enhance total XBP1 (TXBP1) expression. WT Control vs. WT IRI, *p* < 0.05; FXR^-/-^IRI vs. WT IRI, *p* < 0.05. (B) FXR deficiency failed to enhance spliced XBP1 (sXBP1) expression. WT IRI vs. WT Control, *p* < 0.01; and FXR^-/-^IRI vs. WT IRI, *p* < 0.05. (C) FXR deficiency failed to enhance unconventional spliced XBP1 (usXBP1) expression. WT IRI vs. WT Control, *p* < 0.01; FXR^-/-^IRI vs. WT IRI, *p* < 0.05. (D) FXR deficiency failed to increase ATF4 expression. FXR^-/-^IRI vs. WT IRI, *p* < 0.05. (E) FXR deficiency failed to increase GRP94 expression. FXR^-/-^IRI vs. FXR^-/-^Control, *p* < 0.05. (F) Liver IRI reduced EDEM expression in the deficiency of FXR. FXR^-/-^Control vs. FXR^-/-^IRI, *p* < 0.05. (G) FXR deficiency increased CHOP expression. WT Control vs. other groups, *p* < 0.05. (H) The lack of FXR enhanced hepatocyte apoptosis. WT Control vs. WT IRI, *p* < 0.05; FXR^-/-^Control vs. FXR^-/-^IRI, *p* < 0.001, and WT IRI vs. FXR^-/-^IRI, *p* < 0.001.

### Both the lack of FXR and liver IRI altered BA homeostasis

FXR represses transcription of the gene encoding CYP7A1 that is the rate-limiting enzyme in BA synthesis, and thus, the lack of FXR thus resulted in increased circulating total BAs (Fig. 4*A*). BA composition analysis demonstrated that the lack of FXR specifically increased circulating TbMCA (Fig. 4*B*), TCA (Fig. 4*D*) and DCA (Fig. 4*F*) when compared to WT control mice. Liver IRI increased circulating total BAs (Fig.4*A*), TbMCA (Fig. 4*B*), bMCA (Fig. 4*C*), TCA (Fig. 4*D*) and TUDCA (Fig. 4*G*) in WT mice. Although CA and DCA were increased in WT liver IRI mice, there were no statistical differences (*p* > 0.05) (*Fig. 4, E and F*). There were no changes to circulating LCA in any groups (Fig. 4*H*). Given that the mutation of FXR affects BA synthesis and transport (28), and IRI may also impacts liver BA homeostasis, we analyzed total BAs in the liver. As expected, the absence of FXR causes BA accumulation of total BAs in the liver; however, IRI results in decreases of liver total BAs in both WT and FXR^-/-^mice (Fig. 4*I*).

**Fig. 4.**
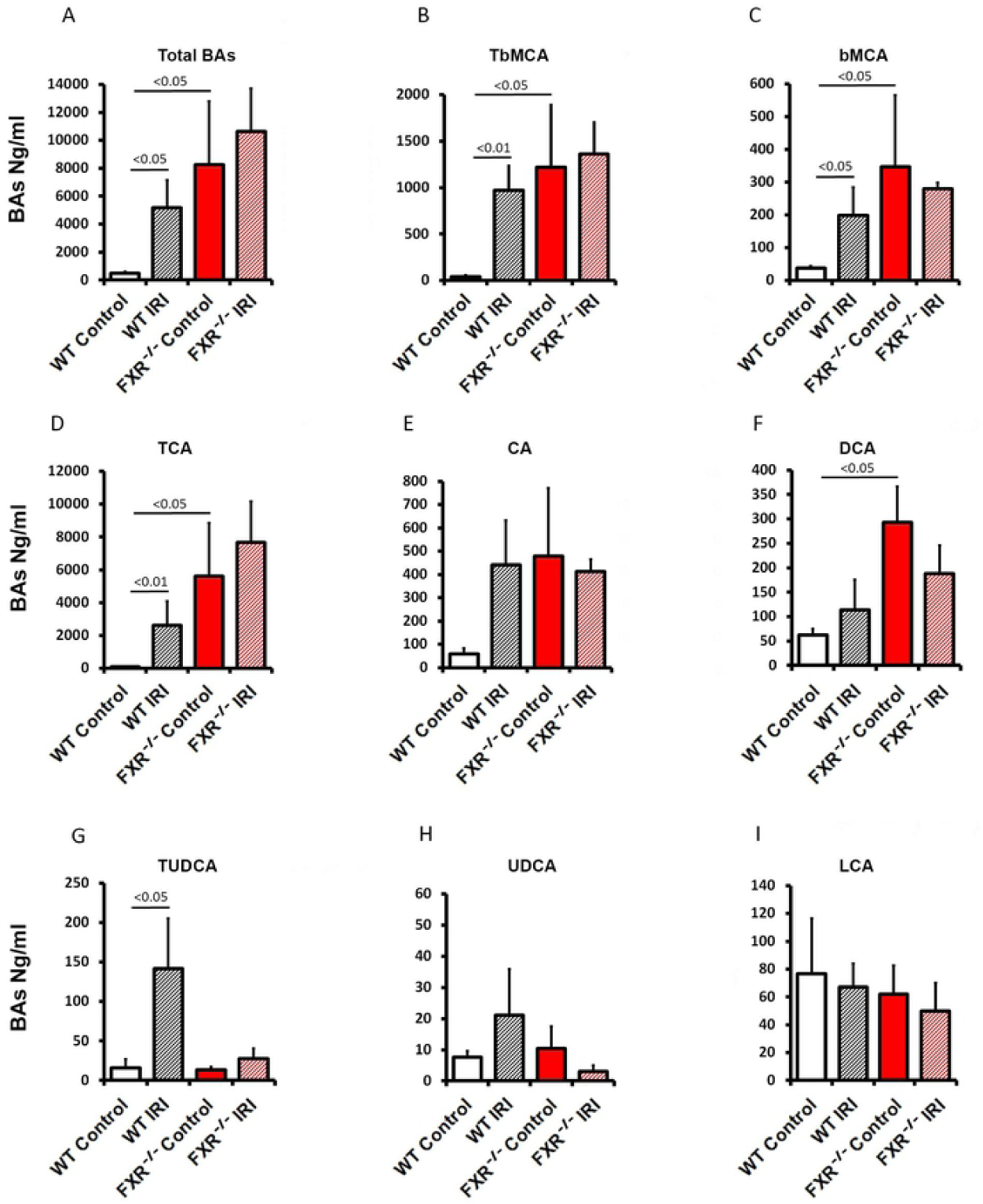
Liver IRI and the lack of FXR altered total BAs and BA composition (n = 5 per group). (A) Liver IRI and the lack of FXR increased circulating total BAs. WT IRI vs. WT Control, *p* < 0.05; FXR^-/-^Control vs. WT Control, *p* < 0.05. (B) Liver IRI and the lack of FXR increased circulating TbMCA. WT IRI vs. WT Control, *p* < 0.01; FXR^-/-^Control vs. WT Control, *p* < 0.05. Liver IRI increased circulating bMCA. WT IRI vs. WT Control, *p* < 0.05; FXR^-/-^Control vs. WT Control, *p* < 0.05. (D) Liver IRI and the lack of FXR increased circulating TCA. WT IRI vs. WT Control, *p* < 0.01; and FXR^-/-^Control vs. WT Control, *p* < 0.05. (E) Liver IRI and the lack of FXR increased circulating CA, but no statistical differences (*p* > 0.05). (F) The lack of FXR increased circulating DCA. FXR^-/-^Control vs. WT Control, *p* < 0.05. (G) Liver IRI increased circulating TUDCA. WT IRI vs. WT Control, *p <* 0.05. (H) Liver IRI increased circulating UDCA in WT mice, but not statistically significant (*p* > 0.05). (I) There were no significant changes of circulating LCA (*p* > 0.05).

Enzyme CYP7A1 initiates the classic pathway of BA synthesis, followed by CYP8B1 that produces the majority of the BA pool. The alternative pathway involves the BA synthetic enzyme CYP27A1 followed by BA hydroxylation by CYP7B1. BA synthesis is under the regulation of FXR, which enhances expression of small heterodimer partner (SHP) and stimulates ileal fibroblast growth factor 15 (FGF15) in mice (FGF19 in humans). FGF15/19 travels to the liver where the membrane receptor, FGFR4, triggers a signaling cascade that results in BA synthesis inactivation. ER stress and inflammatory responses suppress BA synthesis and enhance BA removal from hepatocytes (29). Our results showed that liver IRI inhibited expression of SHP (Fig. 5*A*). Liver IRI and the lack of FXR reduced FGFR4 gene, but no statistical difference (Fig. 5*B*). BA synthesis enzymes, including CYP7A1 (Fig. 5*C*), CYP8B1 (Fig. 5*D*), CYP27A1 (Fig. 5*E*) and CYP7B1 (Fig. 5*F*), were reduced in WT and FXR^-/-^mice. Naïve FXR^-/-^animals demonstrated decreased mean expression levels of CYP27A1 and CYP7B1 mRNAs, but not significantly (*Fig. 5, E and F*, p > 0.05).

**Fig. 5.**
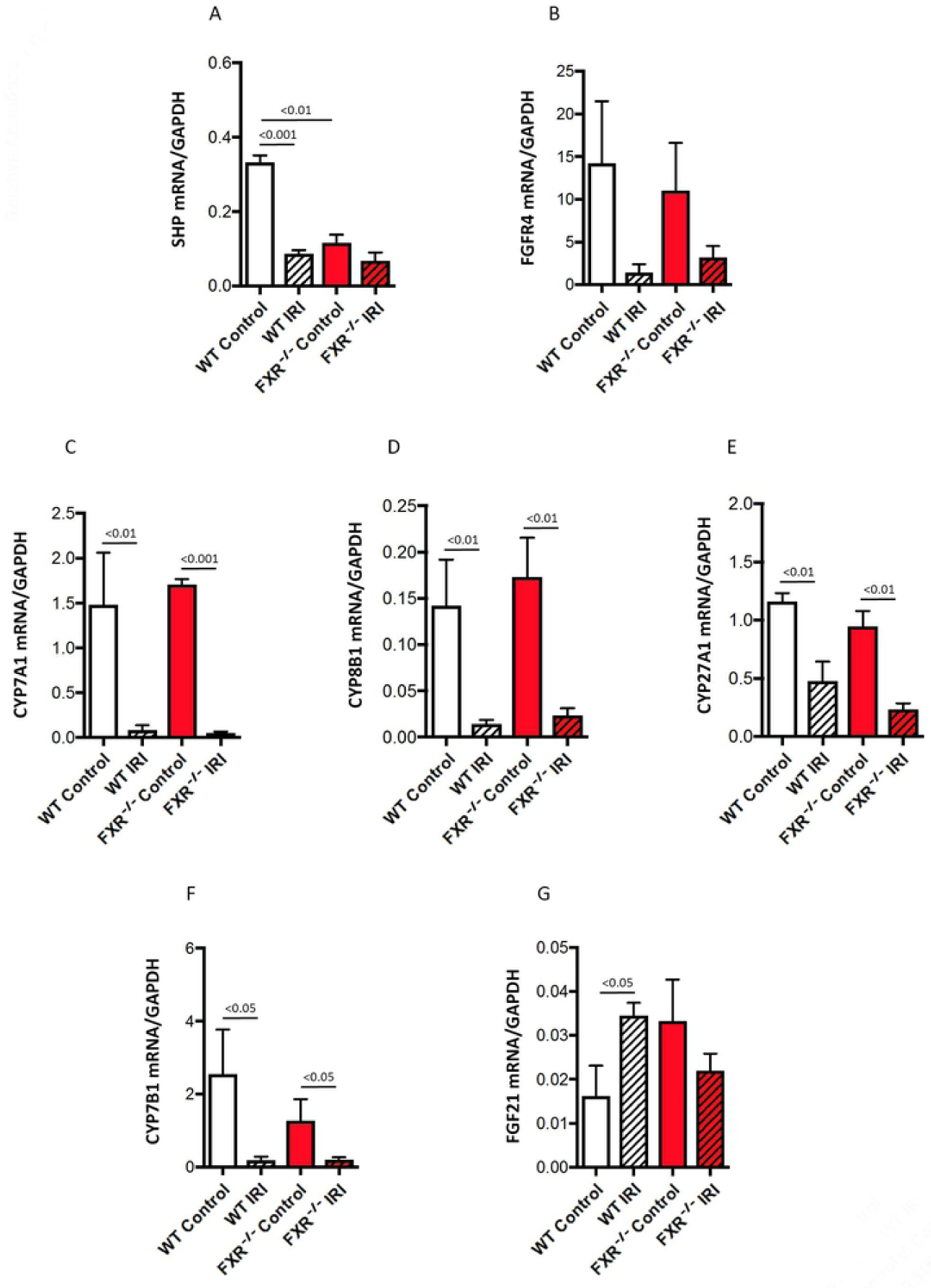
Liver IRI and the lack of FXR altered expression of BA synthesis-related molecules (n = 5 per group). (A) Liver IRI and the lack of FXR decreased expression of SHP in the liver. WT Control vs. WT IRI, *p* < 0.001; FXR^-/-^Control vs. WT Control, *p* < 0.001. (B) Liver IRI decreased expression of FGFR4, but not statistically significant (*p* > 0.05). (C) Liver IRI decreased expression of CYP7A1. WT Control vs. WT IRI, *p* < 0.01; FXR^-/-^IRI vs. FXR^-/-^Control, *p* < 0.001. (D) Liver IRI decreased expression of CYP8B1. WT IRI and FXR^-/-^IRI vs. WT Control and FXR^-/-^Control, *p* < 0.01, respectively. (E) Liver IRI decreased expression of CYP27A1. WT IRI and FXR^-/-^IRI vs. WT Control and FXR^-/-^Control, *p* < 0.01, respectively. (F) Liver IRI decreased expression of CYP7B1. WT Control vs. WT IRI, and FXR^-/-^Control vs. FXR^-/-^IRI, *p* < 0.05, respectively. (G) Liver IRI increased FGF21 expression in WT mice, but not in FXR^-/-^mice. WT Control vs. WT IRI, *p* < 0.05.

Fibroblast growth factor 21 (FGF21) is identified as one of tissue protective proteins (30). To test whether FGF21 is altered under liver IRI, liver FGF21 expression was analyzed. Results showed that liver IRI increased FGF21 transcript in the WT liver. However, IRI failed to enhance liver FGF21 expression in the absence of FXR (Fig. 5*G*), suggesting FGF21 expression may be, at least in part, regulated by FXR (31).

### FXR deficiency altered gut microbiota composition

Taxonomic analysis of gut microbiota composition showed changes at the phylum level, where Bacteroidetes was increased and Firmicutes decreased - with increased Bacteroidetes/Firmicutes ratios - in FXR^-/-^IRI mice with and without liver IRI (*Fig. 6, A and B*) consistent with a previous report (14). Our data showed that Proteobacteria was increased in untreated FXR^-/-^and FXR^-/-^IRI mice (Fig. 6*C*). More interestingly, altered Bacteroidetes and Firmicutes were correlated with markers of liver function; Bacteroidetes was positively and Firmicutes was negatively correlated with serum ALT levels (*Fig. 6, D and E*) following liver IRI. Increasing data identify Proteobacteria as lipopolysaccharide (LPS) producers that act as possible microbial signature of host disease (32). Within the phylum Bacteroidetes, the relative abundance of *Bacteroides* was increased in naïve and FXR^-/-^IRI mice (*Fig. 6, F and G*). Within the phylum Firmicutes, the relative abundances of *Turicibacter* and *Clostridiales* were decreased in FXR^-/-^mice with and without liver IRI (*Fig. 6, H and I*). Finally, within the phylum Proteobacteria, the relative abundance of *Desulfovibrio* was increased in FXR^-/-^mice with and without liver IRI (Fig. 6*J*).

**Fig. 6.**
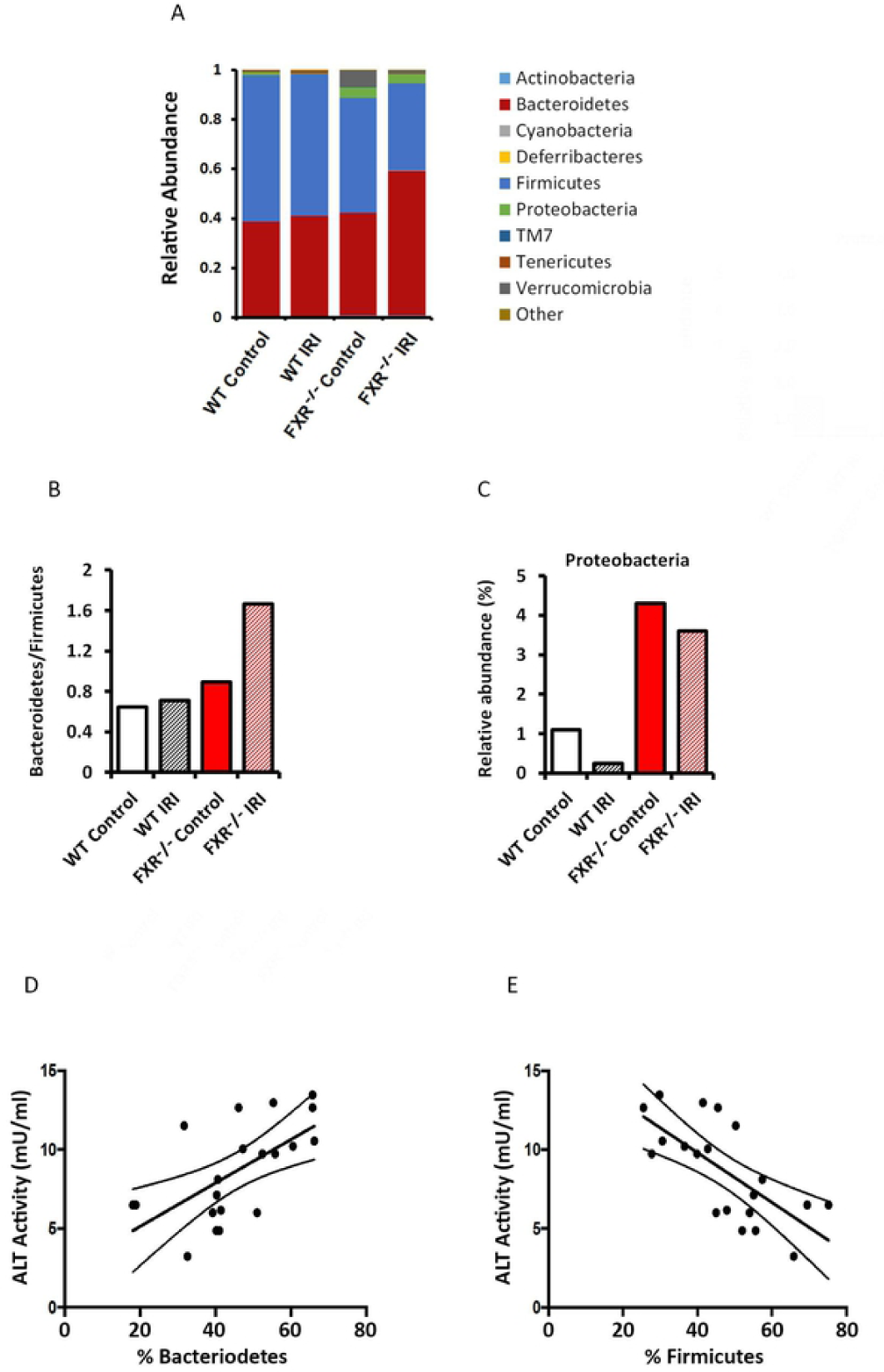
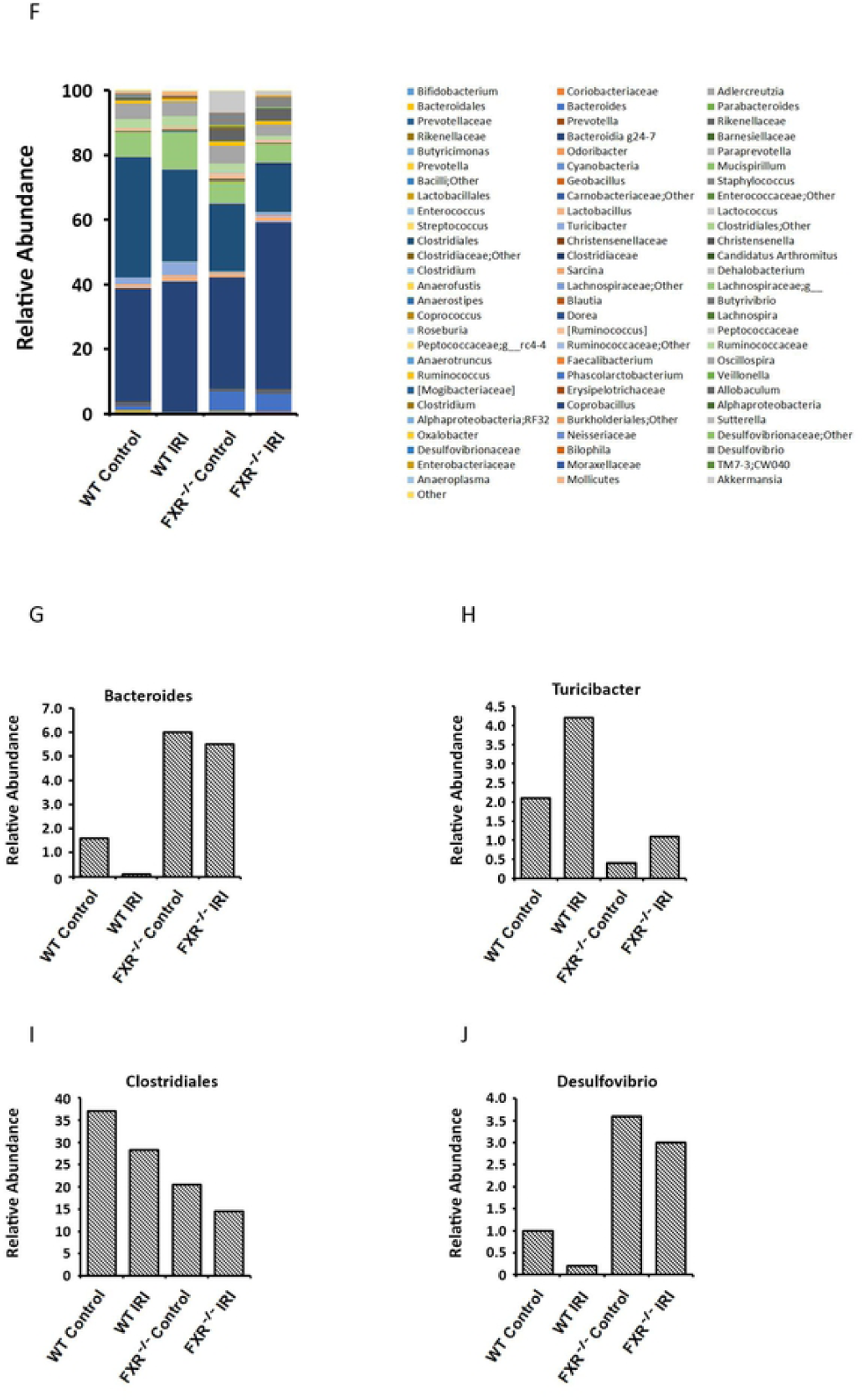
The lack of FXR altered gut microbiota composition (n = 5 per group). (A) Reconfiguration of phyla. The lack of FXR reduced phylum Firmicutes and increased phylum Bacteroidetes. (B) Decreased Firmicutes and increased Bacteroidetes cause an increase in the ratio of Bacteroidetes/Firmicutes in FXR^-/-^mice. (C) The lack of FXR increased phylum Proteobacteria. Increased phylum Bacteroidetes was positively related with increased serum ALT levels in FXR^-/-^mice. (E) Increased phylum Firmicutes was negatively related with serum ALT levels in FXR^-/-^mice. (F) The relative abundance of cecal content genera of microbiome. (G) The lack of FXR led to increase in *Bacteroides* (under phylum Bacteroidetes). (H) The lack of FXR caused decrease of *Turicibacter* (under phylum Firmicutes). (I) The lack of FXR caused decrease of *Clostridiales* (under phylum Firmicutes). (J) The lack of FXR caused increase of *Desulfovibrio* (under phylum Proteobacteria).

**Fig. 7.**
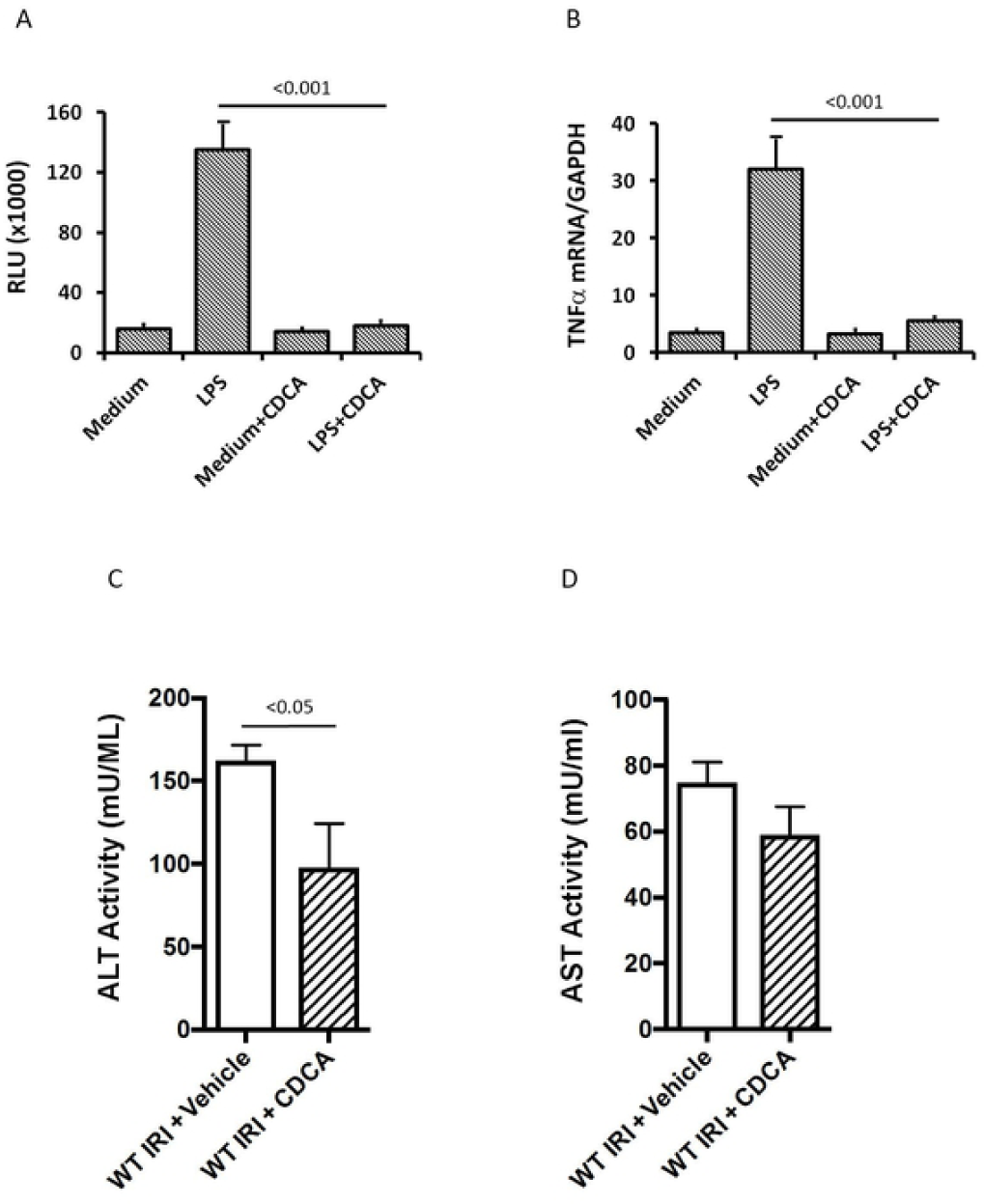

### Activation of FXR inhibited inflammatory reaction *in vitro* and improved liver IRI *in vivo*

Macrophage-induced inflammatory responses are mediated through inflammatory signaling pathways, such as NF-*κ*B, causing TNFα expression. In the bone marrow-derived macrophage (BMDM) culture, LPS increased luciferase activity, indicating increased NF-*κ*B activity, which was inhibited by administration of chenodeoxycholic acid (CDCA, Supplemental Fig. 1S*A*). TNFα mRNA was increased by the stimulation of LPS but also inhibited by CDCA (Supplemental Fig. 1S*B*).

To test whether FXR activation improved liver IRI, FXR agonist CDCA was administered to WT mice undergone 90 minutes of ischemia and 24 hours of reperfusion. The results showed that serum ALT levels were significantly decreased in CDCA-treated WT mice compared to Vehicle-treated WT mice (Supplemental Fig. 1S*C*). Although CDCA also decreased AST levels in the same animals, differences between CDCA-treated and control mice did not reach significance (*p* > 0.05, Supplemental Fig. 1S*D*).

## Discussion

We employed animals lacking FXR compared with WT to investigate critical signaling pathways in liver IRI. The results herein demonstrated that liver IRI decreased hepatic FXR expression in WT mice in line with elevated IL-6 levels, a known FXR repressor (33), indicating an inverse relationship between FXR and liver IRI. Under FXR deficiency, more profound increases in pro-inflammatory cytokines may directly contribute to elevated susceptibility to liver IRI, including elevated circulating AST and ALT, and more pronounced hepatocyte apoptosis compared with WT controls.

ER stress triggers the adaptive response that acts in concert to mitigate the load of new proteins entering the ER through increasing ER content, such as chaperone capacity, thereby degrading misfolded proteins and inhibiting cell apoptosis. Chaperone GRP94 and ATF4 are the most abundant glycoprotein in the ER and participate in protein folding and assist in the targeting of misfolded proteins for ER associated degradation (ERAD) (34). Promotion of ERAD through upregulation of ERAD-enhancing α-mannosidase-like protein (EDEM) promotes cell recovery by accelerating ERAD of terminally misfolded polypeptides and protects against cell apoptosis (35). A recent report suggests that FXR signaling modulates ER stress via activation of the IRE/XBP1 pathway (11). Our findings show that mice lacking FXR resulted in reduced expression of GRP94, ATF4, EDEM, XBP1 (including total, spliced and unconventional spliced XBP1) in the liver when suffering from IRI, suggesting that the absence of FXR forfeits protection of hepatocytes from apoptosis, contributing to the susceptibility to liver IRI.

ROS generation inflicts tissue damage and initiates a cascade of deleterious cellular responses leading to inflammation, cell death and ultimately liver failure. SOD2 transforms toxic superoxide and clears mitochondrial ROS and therefore confers protection against hepatocyte apoptosis (36). In the absence of FXR, liver IRI fails to promote production of SOD2, and accumulation of ROS may initiate the inflammatory reaction and liver damage.

FGF21 is a member of the fibroblast growth factor family that regulates cell growth, differentiation, and glucose and lipid metabolism. FXR activation promotes FGF21 expression (31), which alleviates hepatic ER stress under the physiological condition (37). Increased FGF21 protects against acetaminophen-induced hepatotoxicity by enhancing antioxidant capacity (38). Clinical findings show that serum FGF21 increases following liver or cardiac ischemia and is associated with protective responses (39). We observed that liver IRI promoted expression of liver FGF21 in WT mice, but deficiency of FXR failed to increase FGF21 expression, thereby losing liver protection.

The lack of FXR stimulates BA synthesis, resulting in increased circulating total BAs, TbMCA, MCA, TCA and DCA. Due to the damage of BA transports in the absence of FXR (28), BAs are accumulated in the liver as evidenced by increased total BAs in the liver of FXR^-/-^mice. Liver IRI causes immediate reduction of synthesis as confirmed by decreased BA synthetic enzymes and reduced total BAs in the liver in both WT and FXR^-/-^mice. However, liver IRI induces rapid increases of circulating total BAs and alterations of BA composition in both WT and FXR^-/-^mice. Among the notable changes in circulating BAs was TUDCA, a hydrophilic BA that regulates ER stress and mediates cytoprotective responses during liver IRI (40); however, FXR^-/-^mice failed to demonstrate elevated TUDCA. Nevertheless, whether the alteration of BA composition mediates liver IRI needs further study.

A biochemical link between the gut microbiota and FXR signaling has been demonstrated (41). Taxonomic tree analysis displaying differential taxa in hierarchical layers supports the concept that FXR deficiency is associated with distinct microbiota and liver IRI may generate selection pressure for certain gut bacterial taxa. Recent work shows that FXR^-/-^mice exhibit reduced levels of the phylum Firmicutes and destabilizes the gut microbiota when compared with WT animals (42). Our findings demonstrate that liver IRI decreased the relative composition of the bacterial phylum Bacteroidetes in WT animals, consistent with previous work (43). FXR deficiency leads to increased Bacteroidetes and reduced Firmicutes. Increased Bacteroidetes and reduced Firmicutes were correlated with higher levels of circulating ALT after liver IRI across all animals. This analysis also showed increased cecal Proteobacteria in the absence of FXR. Proteobacteria are a major phylum of gram-negative bacteria that include a wide variety of pathogens, such as *Escherichia Coli*, capable of producing LPS (44). However, Proteobacteria are not increased in WT IRI mice, suggesting that the lack of FXR, but not stress (IRI) contributes to increased Proteobacteria. Within the Proteobacteria, the genus *Desulfovibrio* is a Gram-negative obligate anaerobic that was elevated by IRI. Increased *Desulfovibrio* is associated with inflammation of gut tissues (45) and may mediate inflammatory responses following liver IRI. Due to the presence of LPS in the outer membrane, increased Proteobacteria may induce inflammatory responses in the liver, where LPS and other danger associated molecular patterns reach the liver via the portal vein. Consequently, a lack of FXR results in dysbiosis of the microbiome that are empirically associated with hepatocyte injury and therefore may contribute to the liver IRI pathogenesis.

BMDMs functionally resemble Kupffer cells and are used to study immune responses that mediate liver injury (46). Here we show that FXR agonist CDCA attenuated NF-*κ*B activity and TNFα expression the BMDM cell culture consistent with a previous report (47). When CDCA is administered to WT animals, it improves liver IRI supported by decreased serum ALT levels. However, a major limitation to our work relates to the evidence that mice and humans are inherently different in the homeostasis of BAs, i.e. the human liver synthesize CA and CDCA as FXR agonists, whereas mice synthesize TbMCA that are considered antagonistic towards FXR.

In conclusion, these data illustrate fundamental roles for FXR in the modulation of liver IRI. FXR expression is decreased during liver IRI in WT mice. Mice lacking FXR reduce baseline chaperone expression and exhibit significant changes of BA composition, which potentially triggers IRI-induced hepatocyte apoptosis. The absence of FXR results in dysbiosis of the gut microbiota, which may further exacerbate liver inflammatory reactions during liver IRI. Our findings suggest that hepatocyte protective mechanisms are impaired in the absence of FXR signaling. As proof of this concept, our work shows that activation of FXR with exogenous CDCA inhibits inflammatory responses *in vitro* and improves liver IRI markers *in vivo*.

## Acknowledgments

We would like to acknowledge Digestive Disease Research Core Center at the University of Chicago (DDRCC), Diabetes Research and Training Center at the University of Chicago (DRTC) and the MPMS core at the University of Tennessee Health Science center for assistance with Bile acid measurements.

BA analysis was conducted at the MPMS Facility at UTHSC.

## Author Contribution

JFP and YL contributed to most *in vitro* and *in vivo* experiments; PR and CG contributed to lab analysis; EBC and DPY designed the experiment, and DPY wrote the manuscript. All authors approved the final version.

## Abbreviations

BA: bile acid
CDCA: chenodeoxycholic acid
EDEM: ERAD-enhancing α-mannosidase-like protein
ER: Endoplasmic reticulum
ERAD: ER associated degradation
FXR: farnesoid X receptor
GRP94: glucose-regulated protein 94
IRI: ischemia reperfusion injury
SHP: small heterodimer partner
TUNEL: Terminal deoxynucleotidyl transferase dUTP nick end labeling
XBP1: X box binding protein 1

---

**Supplemental Fig. 1.** FXR agonist chenodeoxycholic acid (CDCA) inhibited inflammatory responses *in vitro* and improved liver function *in vivo*. (A) Supplement of CDCA inhibited luciferase activity induced by LPS in the bone marrow-derived macrophage (BMDM) culture, indicating the inhibition of NF-*κ*B activity. LPS CDCA vs. LPS, *p* < 0.001. (B) Supplement of CDCA inhibited TNFα expression induced by LPS in the BMDM culture. LPS CDCA vs. LPS, *p* < 0.001 (the cell culture was repeated twice). (C) Administration of CDCA decreased serum ALT levels in WT mice with liver IRI (CDCA vs. Vehicle, *p* < 0.05). (D) Serum AST levels in WT mice with liver IRI (CDCA vs. Vehicle, *p* > 0.05).

## References

1. Farges, O., Goutte, N., Bendersky, N., Falissard, B., and Group, A. C.-F. H. S. (2012) Incidence and risks of liver resection: an all-inclusive French nationwide study. Annals of surgery 256, 697–704; discussion 704-695

2. Fayek, S. A., Quintini, C., Chavin, K. D., and Marsh, C. L. (2016) The Current State of Liver Transplantation in the United States: Perspective From American Society of Transplant Surgeons (ASTS) Scientific Studies Committee and Endorsed by ASTS Council. American journal of transplantation: official journal of the American Society of Transplantation and the American Society of Transplant Surgeons 16, 3093–3104

3. Nakamura, K., Kageyama, S., Yue, S., Huang, J., Fujii, T., Ke, B., Sosa, R. A., Reed, E. F., Datta, N., Zarrinpar, A., Busuttil, R. W., and Kupiec-Weglinski, J. W. (2018) Heme oxygenase-1 regulates sirtuin-1-autophagy pathway in liver transplantation: From mouse to human. American journal of transplantation: official journal of the American Society of Transplantation and the American Society of Transplant Surgeons 18, 1110–1121

4. Sugawara, Y., Kubota, K., Ogura, T., Esumi, H., Inoue, K., Takayama, T., and Makuuchi, M. (1998) Increased nitric oxide production in the liver in the perioperative period of partial hepatectomy with Pringle’s maneuver. Journal of hepatology 28, 212–220

5. Melgar-Lesmes, P., Balcells, M., and Edelman, E. R. (2017) Implantation of healthy matrix-embedded endothelial cells rescues dysfunctional endothelium and ischaemic tissue in liver engraftment. Gut 66, 1297–1305

6. Liu, J., Ren, F., Cheng, Q., Bai, L., Shen, X., Gao, F., Busuttil, R. W., Kupiec-Weglinski, J. W., and Zhai, Y. (2012) Endoplasmic reticulum stress modulates liver inflammatory immune response in the pathogenesis of liver ischemia and reperfusion injury. Transplantation 94, 211–217

7. Rao, J., Yue, S., Fu, Y., Zhu, J., Wang, X., Busuttil, R. W., Kupiec-Weglinski, J. W., Lu, L., and Zhai, Y. (2014) ATF6 mediates a pro-inflammatory synergy between ER stress and TLR activation in the pathogenesis of liver ischemia-reperfusion injury. American journal of transplantation: official journal of the American Society of Transplantation and the American Society of Transplant Surgeons 14, 1552–1561

8. Chazouilleres, O., Ballet, F., Legendre, C., Bonnefis, M. T., Rey, C., Chretien, Y., and Poupon, R. (1991) Effect of bile acids on ischemia-reperfusion liver injury. Journal of hepatology 13, 318–322

9. Ejiri, S., Eguchi, Y., Kishida, A., Ishigami, F., Kurumi, Y., Tani, T., and Kodama, M. (2001) Cellular distribution of thrombomodulin as an early marker for warm ischemic liver injury in porcine liver transplantation: protective effect of prostaglandin I2 analogue and tauroursodeoxycholic acid. Transplantation 71, 721–726

10. Yang, H., Zhou, H., Zhuang, L., Auwerx, J., Schoonjans, K., Wang, X., Feng, C., and Lu, L. (2017) Plasma membrane-bound G protein-coupled bile acid receptor attenuates liver ischemia/reperfusion injury via the inhibition of toll-like receptor 4 signaling in mice. Liver transplantation: official publication of the American Association for the Study of Liver Diseases and the International Liver Transplantation Society 23, 63–74

11. Liu, X., Guo, G. L., Kong, B., Hilburn, D. B., Hubchak, S. C., Park, S., LeCuyer, B., Hsieh, A., Wang, L., Fang, D., and Green, R. M. (2018) Farnesoid X receptor signaling activates the hepatic X-box binding protein 1 pathway in vitro and in mice. Hepatology 68, 304–316

12. Ceulemans, L. J., Verbeke, L., Decuypere, J. P., Farre, R., De Hertogh, G., Lenaerts, K., Jochmans, I., Monbaliu, D., Nevens, F., Tack, J., Laleman, W., and Pirenne, J. (2017) Farnesoid X Receptor Activation Attenuates Intestinal Ischemia Reperfusion Injury in Rats. PloS one 12, e0169331

13. Swann, J. R., Want, E. J., Geier, F. M., Spagou, K., Wilson, I. D., Sidaway, J. E., Nicholson, J. K., and Holmes, E. (2011) Systemic gut microbial modulation of bile acid metabolism in host tissue compartments. Proceedings of the National Academy of Sciences of the United States of America 108 Suppl 1, 4523–4530

14. Sheng, L., Jena, P. K., Hu, Y., Liu, H. X., Nagar, N., Kalanetra, K. M., French, S. W., French, S. W., Mills, D. A., and Wan, Y. Y. (2017) Hepatic inflammation caused by dysregulated bile acid synthesis is reversible by butyrate supplementation. The Journal of pathology 243, 431–441

15. Ren, Z., Cui, G., Lu, H., Chen, X., Jiang, J., Liu, H., He, Y., Ding, S., Hu, Z., Wang, W., and Zheng, S. (2013) Liver ischemic preconditioning (IPC) improves intestinal microbiota following liver transplantation in rats through 16s rDNA-based analysis of microbial structure shift. PloS one 8, e75950

16. Abe, Y., Hines, I. N., Zibari, G., Pavlick, K., Gray, L., Kitagawa, Y., and Grisham, M. B. (2009) Mouse model of liver ischemia and reperfusion injury: method for studying reactive oxygen and nitrogen metabolites in vivo. Free radical biology & medicine 46, 1–7

17. Pierre, J. F., Martinez, K. B., Ye, H., Nadimpalli, A., Morton, T. C., Yang, J., Wang, Q., Patno, N., Chang, E. B., and Yin, D. P. (2016) Activation of bile acid signaling improves metabolic phenotypes in high-fat diet-induced obese mice. American journal of physiology. Gastrointestinal and liver physiology 311, G286–304

18. Pierre, J. F., Li, Y., Gomes, C. K., Rao, P., Chang, E. B., and Yin, D. P. (2019) Bile Diversion Improves Metabolic Phenotype Dependent on Farnesoid X Receptor (FXR). Obesity 27, 803–812

19. Caporaso, J. G., Kuczynski, J., Stombaugh, J., Bittinger, K., Bushman, F. D., Costello, E. K., Fierer, N., Pena, A. G., Goodrich, J. K., Gordon, J. I., Huttley, G. A., Kelley, S. T., Knights, D., Koenig, J. E., Ley, R. E., Lozupone, C. A., McDonald, D., Muegge, B. D., Pirrung, M., Reeder, J., Sevinsky, J. R., Turnbaugh, P. J., Walters, W. A., Widmann, J., Yatsunenko, T., Zaneveld, J., and Knight, R. (2010) QIIME allows analysis of high-throughput community sequencing data. Nat Methods 7, 335–336

20. Lee, C. K., Herbold, C. W., Polson, S. W., Wommack, K. E., Williamson, S. J., McDonald, I. R., and Cary, S. C. (2012) Groundtruthing next-gen sequencing for microbial ecology-biases and errors in community structure estimates from PCR amplicon pyrosequencing. PloS one 7, e44224

21. Segata, N., Izard, J., Waldron, L., Gevers, D., Miropolsky, L., Garrett, W. S., and Huttenhower, C. (2011) Metagenomic biomarker discovery and explanation. Genome biology 12, R60

22. Yin, D., Ding, J. W., Shen, J., Ma, L., Hara, M., and Chong, A. S. (2006) Liver ischemia contributes to early islet failure following intraportal transplantation: benefits of liver ischemic-preconditioning. American journal of transplantation: official journal of the American Society of Transplantation and the American Society of Transplant Surgeons 6, 60–68

23. Ma, L. L., Gao, X., Liu, L., Xiang, Z., Blackwell, T. S., Williams, P., Chari, R. S., and Yin, D. P. (2009) CpG oligodeoxynucleotide triggers the liver inflammatory reaction and abrogates spontaneous tolerance. Liver transplantation: official publication of the American Association for the Study of Liver Diseases and the International Liver Transplantation Society 15, 915–923

24. Han, W., Joo, M., Everhart, M. B., Christman, J. W., Yull, F. E., and Blackwell, T. S. (2009) Myeloid cells control termination of lung inflammation through the NF-kappaB pathway. Am J Physiol Lung Cell Mol Physiol 296, L320–327

25. Yan, W., Xu, R., Ma, L. L., Han, W., Geevarghese, S. K., Williams, P. E., Sciammas, R., Chong, A. S., and Yin, D. P. (2013) B cells assist allograft rejection in the deficiency of protein kinase c-theta. Transplant international: official journal of the European Society for Organ Transplantation

26. Chu, M. J., Premkumar, R., Hickey, A. J., Jiang, Y., Delahunt, B., Phillips, A. R., and Bartlett, A. S. (2016) Steatotic livers are susceptible to normothermic ischemia-reperfusion injury from mitochondrial Complex-I dysfunction. World journal of gastroenterology 22, 4673–4684

27. Oslowski, C. M., and Urano, F. (2011) Measuring ER stress and the unfolded protein response using mammalian tissue culture system. Methods in enzymology 490, 71–92

28. Gomez-Ospina, N., Potter, C. J., Xiao, R., Manickam, K., Kim, M. S., Kim, K. H., Shneider, B. L., Picarsic, J. L., Jacobson, T. A., Zhang, J., He, W., Liu, P., Knisely, A. S., Finegold, M. J., Muzny, D. M., Boerwinkle, E., Lupski, J. R., Plon, S. E., Gibbs, R. A., Eng, C. M., Yang, Y., Washington, G. C., Porteus, M. H., Berquist, W. E., Kambham, N., Singh, R. J., Xia, F., Enns, G. M., and Moore, D. D. (2016) Mutations in the nuclear bile acid receptor FXR cause progressive familial intrahepatic cholestasis. Nature communications 7, 10713

29. Henkel, A. S., LeCuyer, B., Olivares, S., and Green, R. M. (2017) Endoplasmic Reticulum Stress Regulates Hepatic Bile Acid Metabolism in Mice. Cellular and molecular gastroenterology and hepatology 3, 261–271

30. Liu, S. Q., Tefft, B. J., Roberts, D. T., Zhang, L. Q., Ren, Y., Li, Y. C., Huang, Y., Zhang, D., Phillips, H. R., and Wu, Y. H. (2012) Cardioprotective proteins upregulated in the liver in response to experimental myocardial ischemia. American journal of physiology. Heart and circulatory physiology 303, H1446–1458

31. Cyphert, H. A., Ge, X., Kohan, A. B., Salati, L. M., Zhang, Y., and Hillgartner, F. B. (2012) Activation of the farnesoid X receptor induces hepatic expression and secretion of fibroblast growth factor The Journal of biological chemistry 287, 25123–25138

32. Carvalho, F. A., Koren, O., Goodrich, J. K., Johansson, M. E., Nalbantoglu, I., Aitken, J. D., Su, Y., Chassaing, B., Walters, W. A., Gonzalez, A., Clemente, J. C., Cullender, T. C., Barnich, N., Darfeuille-Michaud, A., Vijay-Kumar, M., Knight, R., Ley, R. E., and Gewirtz, A. T. (2012) Transient inability to manage proteobacteria promotes chronic gut inflammation in TLR5-deficient mice. Cell host & microbe 12, 139–152

33. Ogura, J., Terada, Y., Tsujimoto, T., Koizumi, T., Kuwayama, K., Maruyama, H., Fujikawa, A., Takaya, A., Kobayashi, M., Itagaki, S., Takahashi, N., Hirano, T., Yamaguchi, H., and Iseki, K. (2012) The decrease in farnesoid X receptor, pregnane X receptor and constitutive androstane receptor in the liver after intestinal ischemia-reperfusion. Journal of pharmacy & pharmaceutical sciences: a publication of the Canadian Society for Pharmaceutical Sciences, Societe canadienne des sciences pharmaceutiques 15, 616–631

34. Zhu, G., and Lee, A. S. (2015) Role of the unfolded protein response, GRP78 and GRP94 in organ homeostasis. Journal of cellular physiology 230, 1413–1420

35. Molinari, M., Calanca, V., Galli, C., Lucca, P., and Paganetti, P. (2003) Role of EDEM in the release of misfolded glycoproteins from the calnexin cycle. Science 299, 1397–1400

36. Pias, E. K., Ekshyyan, O. Y., Rhoads, C. A., Fuseler, J., Harrison, L., and Aw, T. Y. (2003) Differential effects of superoxide dismutase isoform expression on hydroperoxide-induced apoptosis in PC-12 cells. The Journal of biological chemistry 278, 13294–13301

37. Maruyama, R., Shimizu, M., Hashidume, T., Inoue, J., Itoh, N., and Sato, R. (2018) FGF21 Alleviates Hepatic Endoplasmic Reticulum Stress under Physiological Conditions. Journal of nutritional science and vitaminology 64, 200–208

38. Ye, D., Wang, Y., Li, H., Jia, W., Man, K., Lo, C. M., Wang, Y., Lam, K. S., and Xu, A. (2014) Fibroblast growth factor 21 protects against acetaminophen-induced hepatotoxicity by potentiating peroxisome proliferator-activated receptor coactivator protein-1alpha-mediated antioxidant capacity in mice. Hepatology 60, 977–989

39. Liu, S. Q., Roberts, D., Kharitonenkov, A., Zhang, B., Hanson, S. M., Li, Y. C., Zhang, L. Q., and Wu, Y. H. (2013) Endocrine protection of ischemic myocardium by FGF21 from the liver and adipose tissue. Scientific reports 3, 2767

40. Ishigami, F., Naka, S., Takeshita, K., Kurumi, Y., Hanasawa, K., and Tani, T. (2001) Bile salt tauroursodeoxycholic acid modulation of Bax translocation to mitochondria protects the liver from warm ischemia-reperfusion injury in the rat. Transplantation 72, 1803–1807

41. Li, F., Jiang, C., Krausz, K. W., Li, Y., Albert, I., Hao, H., Fabre, K. M., Mitchell, J. B., Patterson, A. D., and Gonzalez, F. J. (2013) Microbiome remodelling leads to inhibition of intestinal farnesoid X receptor signalling and decreased obesity. Nature communications 4, 2384

42. Jena, P. K., Sheng, L., Liu, H. X., Kalanetra, K. M., Mirsoian, A., Murphy, W. J., French, S. W., Krishnan, V. V., Mills, D. A., and Wan, Y. Y. (2017) Western Diet-Induced Dysbiosis in Farnesoid X Receptor Knockout Mice Causes Persistent Hepatic Inflammation after Antibiotic Treatment. The American journal of pathology 187, 1800–1813

43. Nardone, G., Compare, D., Liguori, E., Di Mauro, V., Rocco, A., Barone, M., Napoli, A., Lapi, D., Iovene, M. R., and Colantuoni, A. (2010) Protective effects of Lactobacillus paracasei F19 in a rat model of oxidative and metabolic hepatic injury. American journal of physiology. Gastrointestinal and liver physiology 299, G669–676

44. Mirpuri, J., Raetz, M., Sturge, C. R., Wilhelm, C. L., Benson, A., Savani, R. C., Hooper, L. V., and Yarovinsky, F. (2014) Proteobacteria-specific IgA regulates maturation of the intestinal microbiota. Gut microbes 5, 28–39

45. Lennon, G., Balfe, A., Bambury, N., Lavelle, A., Maguire, A., Docherty, N. G., Coffey, J. C., Winter, D. C., Sheahan, K., and O’Connell, P. R. (2014) Correlations between colonic crypt mucin chemotype, inflammatory grade and Desulfovibrio species in ulcerative colitis. Colorectal disease: the official journal of the Association of Coloproctology of Great Britain and Ireland 16, O161–169

46. Klein, I., Cornejo, J. C., Polakos, N. K., John, B., Wuensch, S. A., Topham, D. J., Pierce, R. H., and Crispe, I. N. (2007) Kupffer cell heterogeneity: functional properties of bone marrow derived and sessile hepatic macrophages. Blood 110, 4077–4085

47. Wang, Y. D., Chen, W. D., Wang, M., Yu, D., Forman, B. M., and Huang, W. (2008) Farnesoid X receptor antagonizes nuclear factor kappaB in hepatic inflammatory response. Hepatology 48, 1632–1643

